# rAAV prostaglandin-based gene therapy lowers intraocular pressure and preserves optic nerve health in glaucomatous DBA/2J mice

**DOI:** 10.64898/2026.03.27.714838

**Authors:** Kristina J. Chern, Rachel L. Fehrman, Gavin J. Marcoe, Daniel M. Lipinski

## Abstract

Open-angle glaucoma (OAG) affects approximately 57.5 million individuals worldwide and is characterized by the progressive loss of retinal ganglion cells (RGC) and irreversible optic nerve damage resulting from chronic ocular hypertension. Intraocular pressure (IOP) is the only major modifiable risk factor in OAG and clinical treatments necessarily aim to lower IOP in order to preserve RGCs and prevent vison loss. Pharmacological therapies, such as prostaglandin analog containing eye drops, are known to be effective at reducing IOP, but are critically undermined by poor patient compliance and are unable to control for potentially damaging diurnal fluctuations in IOP, leading to vision loss even in patient’s diagnosed early. Herein we evaluate the effectiveness of a long-acting, single use, prostaglandin-based recombinant adeno-associated virus (rAAV)-mediated IOP-lowering gene therapy treatment in glaucomatous DBA/2J mice and demonstrate that sustained IOP reduction leads to preservation of both optic nerve anatomy and function in end-stage glaucomatous disease.

**One Sentence Summary:** IOP-lowering gene therapy provides partial anatomical and functional rescue in glaucomatous mouse model following single dose treatment

## INTRODUCTION

Open-angle glaucoma (OAG) affects approximately 57.5 million individuals worldwide and is a leading cause of irreversible blindness characterized by progressive loss of retinal ganglion cells (RGCs) and atrophy of the optic nerve leading to severe visual field defects.[1, 2] Whilst genetics and environment both contribute towards the development and progression of OAG, the only major modifiable risk factor is increased intraocular pressure (IOP), which results from an imbalance between the aqueous humor production from the ciliary body and drainage through the trabecular meshwork.[3] As a consequence, current treatment paradigms for OAG focus almost exclusively on reducing IOP and first-line treatment usually consists of one or more drugs applied topically as eye drops that act to either decrease aqueous humor production (e.g. carbonic anhydrase inhibitors or beta blockers) or promote (e.g. prostaglandin analogs).[4-6] Eye drops containing prostaglandin F_2α_ (PGF_2α_) analogs in particular are extensively prescribed and function to promote drainage through the unconventional pathway via relaxation of the ciliary muscle and remodelling of the surrounding extracellular matrix.[7] Problematically, although topical drug therapies are effective at reducing IOP, they are short-lived, necessitating patients adhere to a daily treatment regimen throughout their lifetime, which negatively affects quality of life and imposes a substantial financial and medical burden.[8] Long-term use of PGF_2α_ analog eye drops is also associated with numerous common side effects, including ocular irritation and hyperemia, darkening of iris pigmentation, abnormal eyelash growth and increased risk of infectious keratitis/conjunctivitis.[9] Owing to these factors and the absence of immediate or noticeable relief following drop application, compliance rates for existing drug-based therapies for OAG are extremely poor, with <25% of patients maintaining treatment over even a 1-year period.[10, 11] Furthermore, drug administration via eye drops necessarily results in periodic bolus dosing and so does not control for natural diurnal fluctuations in IOP – an independent risk factor for OAG progression – and so may fail to prevent glaucomatous vision loss even in patients diagnosed early and who comply with therapy. [12, 13]

Owing to the limitations of existing pharmacological approaches to lower IOP there has been a drive to develop longer acting therapies to treat OAG, including slow-release formulations of existing drugs, such as brimonidine or bimatoprost, delivered from either a sustained release injectable implant or topical solid polymer.[14, 15] Whilst the most advanced of these technologies (e.g. Durysta intracameral implant) have been successful in achieving a clinically relevant reduction in IOP in human subjects for a period of up to several months, their impact is likely to be substantially limited by the potential for serious side adverse events, including macular edema, inflammation and corneal endothelial cell loss, which together contraindicate implant re-administration – a major limitation for the treatment of a chronic lifelong condition.[16-19]

More recently, alternative approaches aimed at vectorizing existing pharmacological treatments using gene therapy to more permanently modulate ‘pathway biology’ have gained traction particularly within the ophthalmology space, where for example, several clinical trials are ongoing to assess whether recombinant adeno-associated virus (rAAV)-mediated expression of vascular endothelial growth factor inhibitors limits neovascularization in wet age-related macular degeneration.[20] Our group and others have similarly explored whether OAG can be treated using a gene therapy approach and have demonstrated that virally vector-mediated *de novo* biosynthesis of native PGF_2α_ can mediate long-term, dose-dependent IOP reduction following a single therapeutic intervention in a variety of normotensive animal species, including rats, cats and non-human primates.[21-23] Whilst highly promising, to date it has not been demonstrated that such a gene therapy approach is effective at lowering IOP in a glaucomatous animal model or whether doing so leads to anatomical and functional preservation of the optic nerve; both prerequisites for clinical translation to human subjects.

In this study, we evaluate the long-term therapeutic effectiveness of rAAV-mediated IOP reduction in the extensively characterized DBA/2J mouse model of congenital glaucoma, which has recessive mutations in the glycoprotein nmb (*Gpnmb*) and Tyrosinase related protein 1 (*Tyrp1*) genes that cause progressive iris stromal atrophy and pigment dispersion leading to obstruction of the trabecular meshwork and IOP elevation.[24, 25] Importantly, although the DBA/2J mouse is a model of secondary glaucoma, it accurately recapitulates several key aspects of human OAG, including optic nerve excavation and RGC loss as a results of chronic ocular hypertension and a responsiveness to prostaglandin analog eye drop treatment, that make then ideally suited for the evaluation of our gene therapy treatment.[23, 24, 26-28]

Using a combination of repetitive rebound tonometry, electroretinography (ERG) and post-mortem histology, herein we demonstrate that a single dose rAAV-mediated gene therapy treatment that catalyzes the *de novo* biosynthesis of native PGF2α and its cognate receptor effectively lowers IOP in the DBA/2J mouse model and leads to partial anatomical and functional rescue. If translated, this IOP-lowering therapy would be expected to have a profound benefit for the clinical management of OAG by allowing for the diurnal maintenance of normal tension while eliminating the necessity for adherence to a daily treatment regimen.

## RESULTS

### Intracameral administration of rAAV2/5.CCPP effectively prevents IOP elevation in DBA/2J mice relative to natural history and HBSS sham treated controls

DBA/2J mice (N=80; 50:50 male/female) were imported from a commercial breeder at 2-months of age and allowed to acclimate to local housing conditions before being assigned randomly to one of four cohorts: untreated natural history control, HBSS sham injected control, or injection with a rAAV2/5 vector packaging the smCBA-TC40-COX2-P2A-PTGFR-TC45 transgene cassette (herein known as rAAV2/5.CCPP), which expresses the rate limiting enzyme (COX2) in the *de novo* biosynthesis of PGF_2α_ and the primary receptor (PTGFR) under control of a ubiquitous promoter, at either a low (4x10^9^ vector genomes (vg)/eye) or high (4x10^10^ vg/eye) dose.[23]

Rebound tonometry was performed bilaterally on isoflurane anesthetized DBA/2J mice (N=160 eyes, 9 averaged recordings per measurement) at baseline (3 months; pre-treatment), 4-months (time of treatment), and 6-, 7-, 8-, 9-, and 10-months (all post-treatment) of age by examiners who were masked with respect to treatment group (Fig. 1A). In line with previously published data, IOP in the DBA/2J mouse was observed to slightly, but non-significantly decrease between 3- and 7-months in all groups (Figure 1B). [26] At 8 months of age, the natural history group demonstrated a highly significant increase in IOP versus both low dose (IOP= 15.70 vs 10.79 mm Hg, +31.25%, p=<0.0001) and high dose (IOP= 15.70 vs 12.31 mm Hg, +21.63%, p=<0.0001) rAAV2/5.CCPP treated groups (two-way ANOVA with Tukey test for multiple comparisons). IOPs in the sham HBSS treated group were also significantly increased at 8-months versus both low dose (IOP= 14.58 vs 10.79 mm Hg, +25.97%, p=<0.0001) and high dose (IOP= 14.58 vs 12.31 mm Hg, +15.61%, p=<0.001) rAAV2/5.CCPP treated groups, indicating robust protection against ocular hypertension following gene therapy treatment.

**Figure 1.**
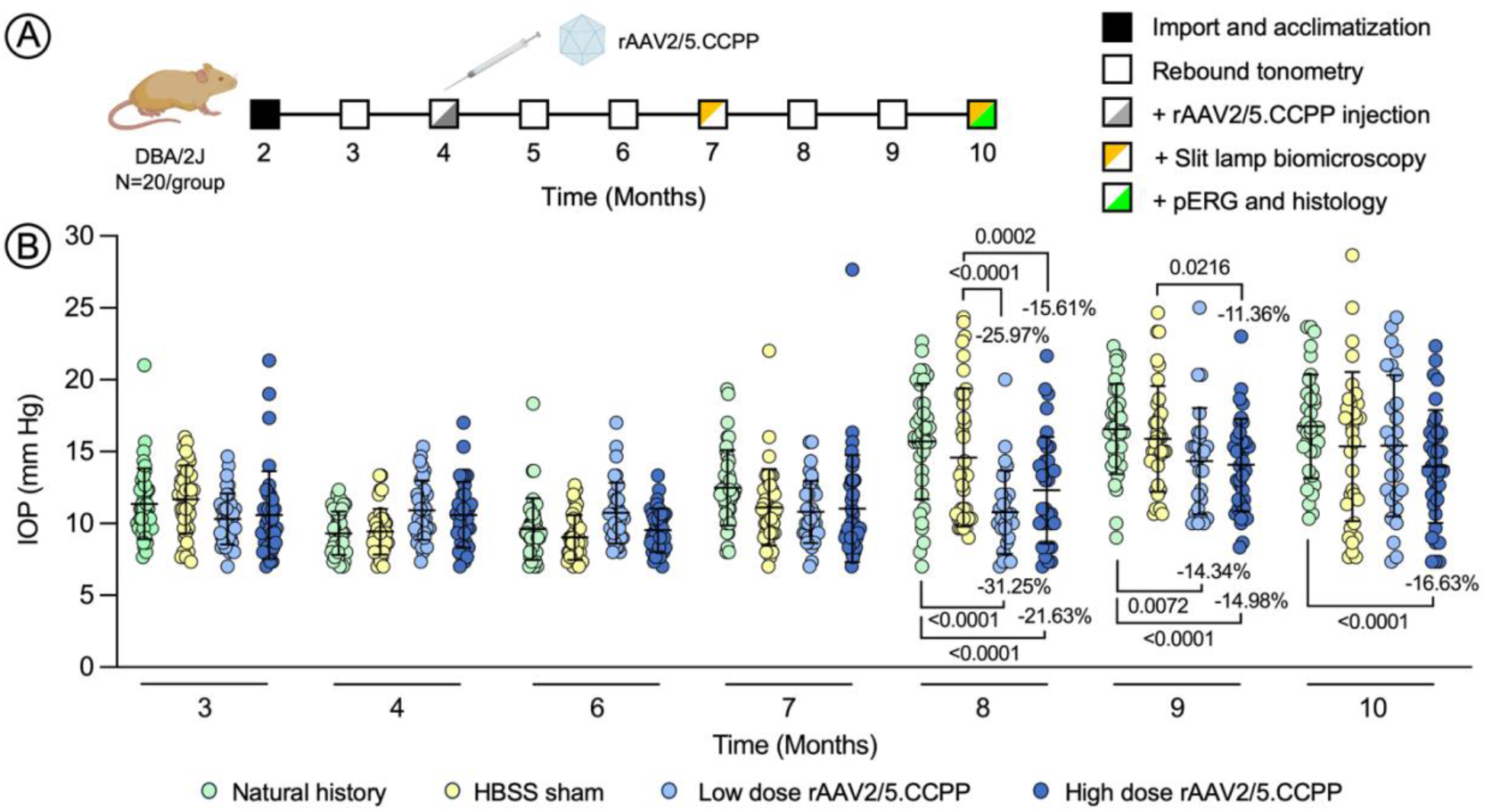
Experimental design and IOP measurements following rAAV2/5.CCPP therapy. Experimental design schematic showing timings of interventions and assessments in DBA/2J mice (**A**) Rebound tonometry measurements of intraocular pressure for natural history control (green), HBSS injected control (yellow), low dose rAAV2/5.CCPP injected (light blue) and high dose rAAV2/5.CCPP injected (dark blue) DBA/2J mice (**B**) Each circle represents the mean of three separate rebound tonometry readings with mean and standard deviation shown. All comparisons two-way ANOVA with Tukey post-test.

IOP in the natural history (16.56 mm Hg) and HBSS sham (15.89 mm Hg) treated groups continued to be significantly elevated (+14.34%, p=<0.01 and +14.98%, p=0.0001, respectively) compared to the low dose rAAV2/5.CCPP treated group (14.35 mm Hg) at 9-months, while IOP in the HBSS sham treated group was also elevated (+11.36%, p=<0.05) relative to the high dose rAAV2/5.CCPP treated eyes (14.08 mm Hg).

At 10-months, IOP remained significantly decreased only in high dose rAAV2/5.CCPP treated eyes relative to the natural history control group (16.75 vs 13.96 mm Hg, +16.63%, p=<0.0001), indicating that rAAV2/5.CCPP treatment remained only partially effective at reducing IOP in end-stage disease, likely owing to a combination of extensive pigment dispersion from the iris causing blockage of the trabecular meshwork that cannot be fully compensated through PGF_2α_-mediated increase in uveoscleral aqueous drainage, and the known reduced sensitivity of DBA/2J mice to IOP reduction with prostaglandins as a function of advancing age.[29]

Slit lamp biomicroscopy performed at 7- and 10-months revealed extensive transillumination defects resulting from stromal atrophy that were comparable across all groups irrespective of treatment (Supplemental Fig. 1) and an absence inflammation (e.g., cell, flare, fibrin strands etc.), strongly indicating that rAAV2/5.CCPP therapy is well tolerated and that the observed reduction in IOP is not the result of chronic anterior uveitis.

### rAAV2/5.CCPP gene therapy partially preserves retinal ganglion cell function

To assess whether rAAV2/5.CCPP treatment led to preservation of visual function at the 10-month timepoint, pattern electroretinography (pERG) was used to measure RGC activity independent of photoreceptor activation. Using a low frequency (1Hz), square-wave pattern reversal stimuli with 100% contrast (black/white) presented under constant luminance (50 cd/m^2^) we were able in the majority of mice to identify the initial negative N_35_ deflection (Figure 3A, grey arrow) and subsequent positive P_50_ deflection (Figure 3A, white arrow), allowing N_35_-P_50_ amplitude to be calculated, which in mice (lacking macular cones) corresponds exclusively to RGC activation. pERG amplitudes recorded unilaterally from natural history (0.81±1.07 μV) and HBSS sham treated (0.49±0.48 μV) groups (N=14 eyes per group) were undetectable or at near noise levels in all animals, indicating wide-spread dysfunction or loss of RGCs resulting from sustained ocular hypertension (Figure 2A-B). By contrast, pERG amplitudes recorded from low (2.53±1.47 μV, N=13 eyes) and high (3.05±2.49 μV, N=14 eyes) dose rAAV2/5.CCPP injected eyes were significantly higher than both natural history (P=<0.05 and <0.01) and HBSS sham treated (P=<0.01 and <0.001) groups, respectively (one-way ANOVA with Tukey posttest), indicating partial protection of RGC function due to rAAV2/5.CCP-mediated IOP reduction in end-stage (10-months) disease.

**Figure 2.**
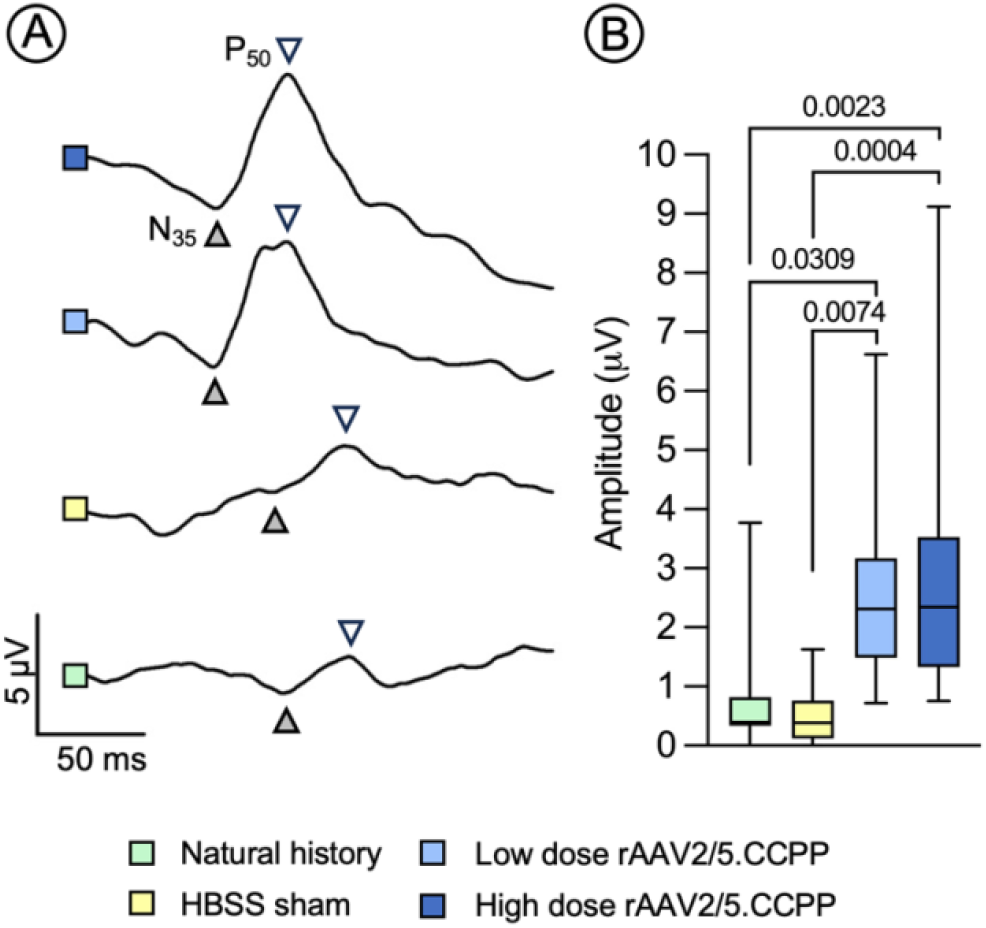
Functional rescue of RGC activity in response to rAAV2/5.CCPP therapy. Representative smoothed pERG traces (**A**) and averaged amplitudes (**B**) at 10-month time point from natural history control (green), HBSS injected control (yellow), low dose rAAV2/5.CCPP injected (light blue) and high dose rAAV2/5.CCPP injected (dark blue) DBA/2J mice demonstrating a statistically significant difference between control and rAAV-treated groups. Grey arrow = N_35_ negative waveform; white arrow = P_50_ positive waveform. Comparisons performed by one-way ANOVA with Tukey post-test.

**Figure 3.**
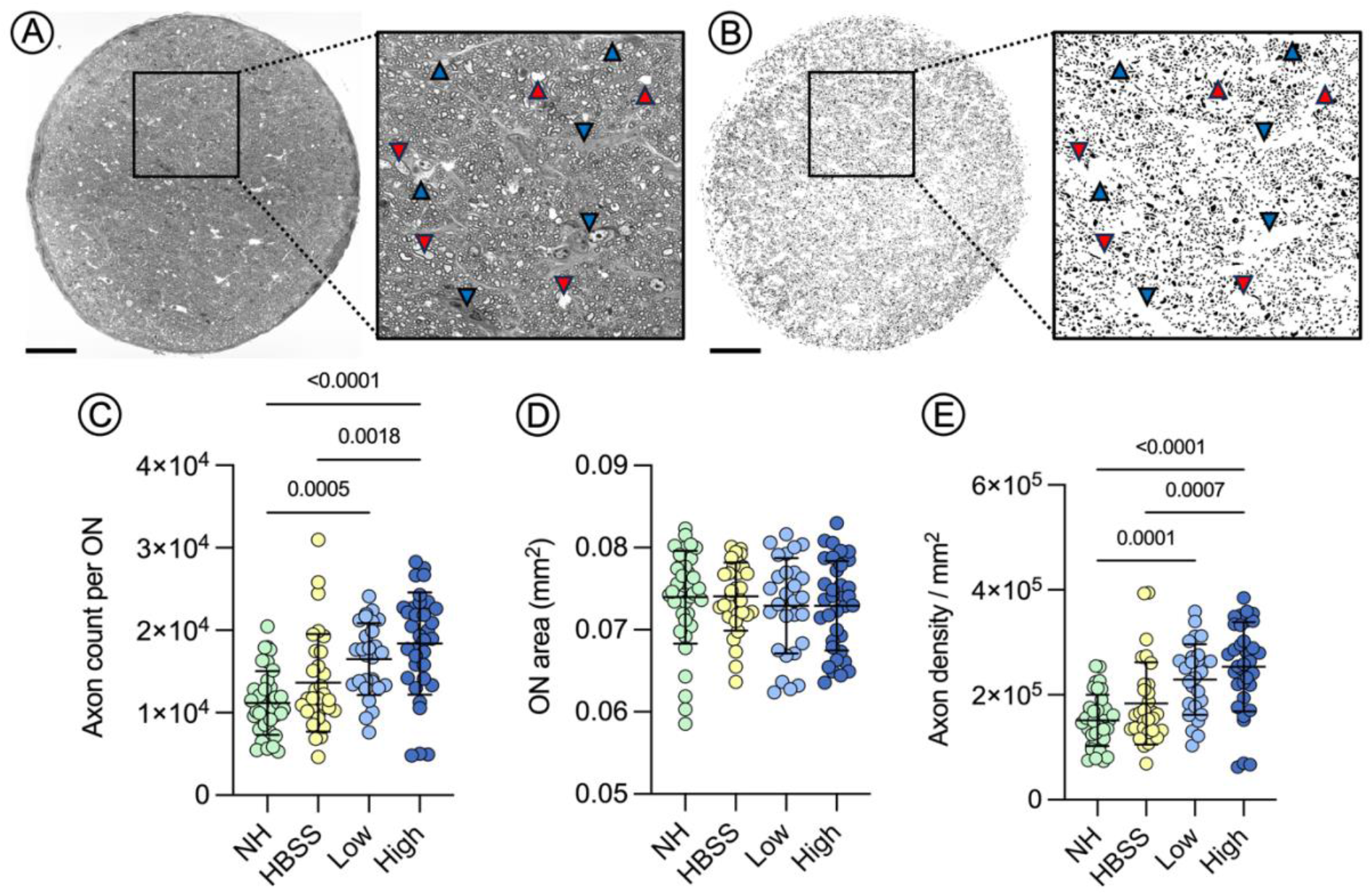
Robust preservation of RGC axon number and density following rAAV2/5.CCPP treatment in 10-month DBA/2J mice. (**A**) Representative light micrograph of a toluidine blue stained coronal optic nerve section showing abundant intact myelinated axons (small white circles with complete black border) distributed amongst areas of axonal swelling (red arrows) and gliosis (blue arrows). (B) Example output from semi-automated quantification approach demonstrating mask of intact axons following exclusion of areas corresponding to swelling (red arrows) and gliosis (blue arrows) based on size and sphericity. Quantification of absolute axon number (**C**) total optic nerve area (**D**) and axon density (**E**) demonstrating robust protection in low dose rAAV2/5.CCPP injected (light blue) and high dose rAAV2/5.CCPP injected (dark blue) compared to natural history control (green) and HBSS injected (yellow) controls. All comparisons performed by one-way ANOVA with Tukey post-test. Scale bar = 50 um

### rAAV2/5.CCPP therapy leads to robust preservation of ganglion cell axons

At 10-months of age – the time point at which IOP elevation in DBA/2J mice has peaked and sustained ocular hypertension is expected to have caused substantial RGC loss and damage to the optic nerve including disc cupping (Supplemental Figure 2) – eyes from animals in all groups were harvested for post-mortem histology in order to assess optic nerve health. Specifically, globes were carefully enucleated so as to preserve ∼2mm of the optic nerve immediately proximal to the sclera before the nerves fixed in formaldehyde-glutaraldehyde, embedded in plastic for semi-thin (6*μm*) sectioning, stained with toluidine blue and imaged via electron microscopy to reveal axonal myelination and ultrastructural anatomy of the nerve. Coronal optic nerve sections harvested from natural history control (N=36), HBSS sham control (N=31), low dose rAAV2/5.CCPP (N=29) and high dose rAAV2/5.CCPP (N=35) injected mice were randomized and analyzed using a semi-automated pipeline that masks areas of axonal swelling/degeneration (Figure 3A-B, red arrows) and gliosis (Figure 3A-B, blue arrows) in order to delineate only intact (i.e. fully myelinated) axons (Figure 3A-B, black dots), allowing for accurate quantification across the entirety of each optic nerve section. Absolute axon quantification per ON section revealed a significantly greater number of intact axons in low dose rAAV2.5.CCPP treated eyes than NH controls (16,497±4,344 vs. 11,183±3,877; P=0.0005), and in high dose rAAV2/5.CCPP treated eyes (18,397±6,218) versus both NH (P=<0.0001) and HBSS (13,634±5,906; P=0.0018) control eyes, respectively (one-way ANOVA with Tukey posttest; Figure 3C).

The cross-sectional area of each optic nerve was not observed to be different between groups (mean = 0.073±0.005mm^2^), indicating an absence of optic nerve atrophy despite differential amounts of axonal loss, swelling and gliosis between eyes (Figure 3D). Consequently, axonal density per ON was also significantly greater in low dose rAAV2.5.CCPP treated eyes than NH controls (228,960±67,359 vs. 151,196±48,973 axons/mm^2^; P=0.0001), and in high dose rAAV2/5.CCPP treated eyes (253,368±85,042 axons/mm^2^) versus both NH (P=<0.0001) and HBSS (182,293±78,380 axons/mm^2^; P=0.007) control eyes, respectively (one-way ANOVA with Tukey posttest; Figure 3C).

### rAAV2/5.CCPP therapy does not protect against gliosis or swelling of the optic nerve

In contrast to axon survival, it is not possible to objectively quantify the extent of optic nerve swelling or gliosis using an automated approach and so five independent graders were presented with scanned light micrographs of each ON section in a masked fashion and asked to score the extent of both swelling and gliosis using a four-point scale ranging from no damage (grade zero) to severe, widespread lesions for each metric (Figure 4A; Supplemental Figure 3 and Supplemental Table 1). Mean scores aggregated across all examiners revealed no significant difference (P>0.05) in the distribution of lesion severity for either swelling or gliosis irrespective of treatment group (Chi squared test), indicating that rAAV-mediated IOP reduction does not impact the extent of inflammation or scarring within the optic nerve.

**Figure 4.**
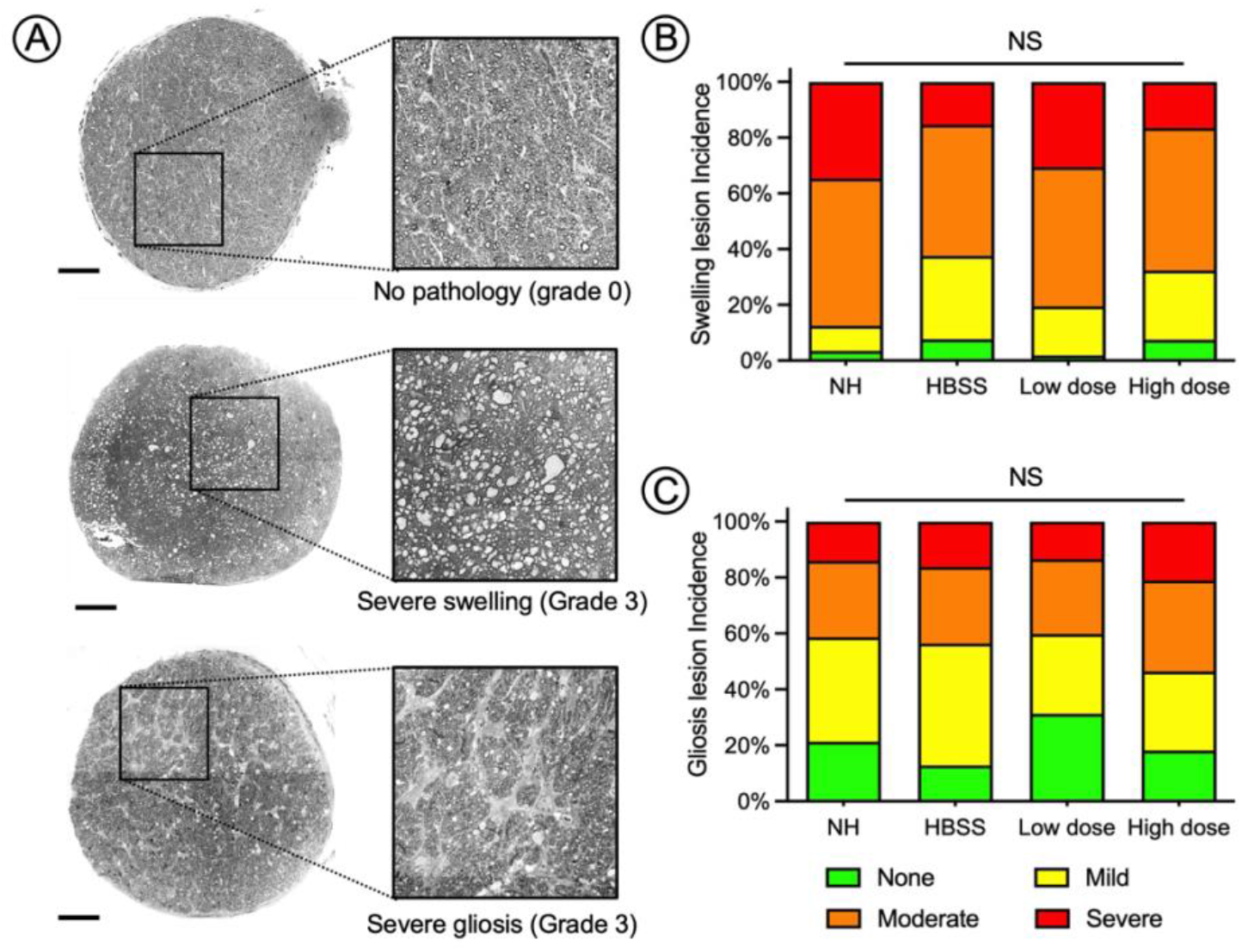
Optic nerve swelling and gliosis are unchanged following rAAV2/5.CCPP treatment at 10-months. (**A**) Representative light micrographs of toluidine blue stained coronal optic nerve sections showing a normal nerve without obvious pathology (top), severe swelling (middle) or severe gliosis (bottom). Stacked bar diagrams showing the distribution of lesion severity for swelling (B) and gliosis (C) across each treatment group as assessed by N=5 independent graders masked to treatment group. Chi-squared testing revealed no significant difference in the proportion of grade severity as a result of rAAV2/5.CCPP treatment. Scale bar = 50 um

## DISCUSSION

Since their identification in the 1980s as potent modulators of IOP in both pre-clinical animal studies and human clinical trials, prostaglandins (e.g. PGF_2α_ and PGE_2_) formulated as esterified pro-drugs and applied topically as eye drops have revolutionized the clinical management of glaucoma and are now the first-line treatment to combat ocular hypertension. [30-33] Unfortunately, while the benefits of using prostaglandin-analog eye drops to reduce IOP in hypertensive patients, particularly those with open-angle glaucoma, has been thoroughly demonstrated in clinical practice, and despite two decades of advances in medicinal chemistry to improve tolerability (i.e. reduce side-effects) and therapeutic efficacy, compliance with prostaglandin analog eye drops remains extremely poor.[10, 11, 34] When combined with the fundamental inability of short-acting topical IOP lowering therapies to compensate for diurnal IOP fluctuations, even in fully compliant patients, there has been a growing drive towards the development of technologies to lower IOP consistently over extended periods.[12, 13] These have primarily taken the form of slow-release formulations of existing pharmacological products, such as brimonidine or bimatoprost, administered either peri- or intra-ocularly as drug-loaded nanogels or solid polymers, and have shown the ability to lower IOP over a period of weeks or months.[14, 15] Although encouraging, the longest acting implantable formulations are associated with substantial side-effects – such as corneal endothelial atrophy and foreign body sensation *in addition* to those observed with traditional topical therapies – that substantially limit their clinical impact by preventing administration on more than one occasion.[19, 35]

In this study, we build upon ours and others previous works and present an alternative, single injection gene therapy approach that functions to reduce IOP by catalyzing the *de novo* biosynthesize and secretion of native PGF_2α_ directly into the anterior chamber.[21-23] Importantly, we demonstrate that that gene therapy can be used to reduce IOP in a gene agnostic manner in a glaucomatous animal model leading to a significant positive effect on both anatomical and functional health of the optic nerve.

As with all animal models of glaucoma, the DBA/2J mouse used throughout this study has both major benefits and limitations. Most importantly and in spite of being a congenital model of pigment dispersion glaucoma, DBA/2J mice accurately recapitulate several aspects of human open-angle glaucoma, including exhibiting progressive optic disc cupping, axonal loss that arises directly from sustained ocular hypertension, and a responsiveness to topical prostaglandin eye drop therapies.[26, 28] In this respect, the model is well suited to evaluate whether vectorizing PGF_2α_ expression is an effective therapeutic strategy and allowed us to comprehensively demonstrate that a clinically relevant ‘target’ reduction in IOP of 20-30% at 8-9 months can be achieved following a single gene therapy intervention (via rebound tonometry; Figure 1), leading to a highly significant (partial) amelioration of both RGC function (assessed via pERG; Figure 2) and optic nerve health (quantified on histology via direct axon counts; Figure 3).[36]

Unfortunately, the particular kinetics of glaucoma progression in the DBA/2J mouse – wherein IOP declines slightly at 3-4 months, increases rapidly around 8-months and then declines precipitously again between 10-12 months as the iridocorneal angle deteriorates – necessitated that mice were pre-treated *before* the onset of glaucoma. As a consequence, we were unable to design our study to address the most clinically relevant question of whether rAAV-mediated PGF_2α_ expression is sufficient to lower IOP to a clinically relevant level in an already glaucomatous eye – a key question that will require further pre-clinical studies in a larger, more slowly progressing model of glaucoma to address, such as the naturally occurring ADAMTS10 beagle model, which has a similar glaucoma phenotype to human subjects and has been used to screen PGF_2α_ based drug treatments previously.[37-39]

Another common feature of the DBA/2J mouse model is the development of corneal neovascularization and calcifications, which affected our ability to resolve the retina using optical coherence tomography (OCT) with sufficient resolution in order to accurately segment individual retinal layers, such as the nerve fiber or ganglion cell layers, which would have allowed for the longitudinal measurement of retinal thinning or optic disc cupping in response to sustained ocular hypertension. It has also been proposed that the corneal phenotype of the DBA/2J mouse may also confound measurement of IOP via rebound tonometry, where calcified deposits affect how the probe rebounds giving unreliable or falsely high recordings.[40] However, as our study protocol involved a very large number of animals, averaged multiple IOP recordings from each eye per time point, and did not observe any systematic groupwise difference in corneal neovascularization or calcification severity that might have influenced data collection, we do not believe that this phenotype had any substantiative effect on the study outcome or conclusion.

One of the major findings of our study is that IOP-reduction in DBA/2J mice following gene therapy treatment led to a preservation of N_35_-P_50_ pERG amplitude in rAAV2/5.CCPP treated eyes (Fig. 2). This is an critical observation as it strongly indicates that RGCs remaining at 10-months of age were functional and likely capable of relaying visually useful information to the brain, although we did not make visually evoked potential (VEP) recordings or other measurements of cortical function (e.g. laser speckle contrast imaging) as part of this study for confirmation.[41, 42] Importantly, pERG amplitudes in both low and high dose rAAV2/5.CCPP treated eyes were significantly higher than both natural history and HBSS sham treated control groups, and were substantially above noise levels reported previously when pERG was performed in similarly aged, untreated DBA/2J mice, strongly indicating that rAAV2/5.CCPP mediated biosynthesis of native PGF_2α_ is sufficient to at least partially rescue the functional phenotype in this model.[43, 44] Unfortunately, to the authors’ knowledge, pERG has not been performed at a comparable time point in DBA/2J mice following long-term (i.e. daily over several months) treatment with PGF_2α_ analog eye drops and so we are unable to comment whether our gene therapy approach is more or less effective at preserving RGC function than traditional therapeutic approaches, or whether there is a limit to the degree of preservation that can be achieved using an IOP-lowering approach.

These findings agree well with histopathological observations of optic nerve health, wherein both low and high dose rAAV2/5.CCPP treatment resulted in a highly significant (P=0.0018 to <0.0001) preservation of intact RGC axons compared to NH and HBSS control groups (Figure 3), strongly indicating that the observed functional preservation is directly correlated to the number of intact RGC axons. Whilst highly encouraging, both absolute axon count per ON (high dose = 18,397±6,218) and axon density (253,368±85,042 axons/mm^2^) observed in rAAV2/5.CCPP treated mice was appreciably lower (∼2-fold) than values reported previously for wild-type non-glaucomatous animals with the same congenic background (e.g. C57BL/6), indicating that the anatomical phenotypic rescue is only partial as a result of rAAV-mediated IOP reduction.[45, 46]

In contrast, the incidence of swelling and gliosis was observed to be unchanged across all groups, with no obvious evidence of rAAV2/5.CCPP treatment increasing or decreasing lesion severity. While every effort was made to ensure the consistency and accuracy of optic nerve grading – with multiple (N=5) independent graders who were masked to treatment group performing the analysis of lesion severity – such assessments inevitably have a high degree of subjectivity. In order to assess the extent to which grading variability may affect our interpretation of the data within each group several images (NH = 6, HBSS = 4, low dose rAAV2/5.CCPP = 4 and high dose rAAV2/5.CCPP = 4) were duplicated, transformed (e.g. rotated and/or flipped) and presented as independent images to each examiner without their knowledge to enable inter-rater reproducibility to be plotted. Grading reproducibility was found to be only fair (*K*=0.370; P=<0.001) for gliosis and moderate for swelling (*K*=0.520; P=<0.001; weighted Cohen’s Kappa; Supplemental Fig. 4). As such, it is unclear whether rAAV2/5.CCPP treatment truly has no effect on the incidence of swelling and gliosis, or whether a small treatment effect (positive or negative) is being obscured by a high degree of grading inconsistency.

In summary, the study presented here provides strong evidence that rAAV-mediated IOP reduction may be a promising therapeutic approach to mitigate both the anatomical and functional manifestations of glaucoma in a manner that could be applied irrespective of the underlying genetic etiology. Importantly, IOP reduction was observed following only a single vector dose and may be maintained long-term – in line with our previous studies conducted in normotensive rats – providing a clear potential advantage over the current clinical gold standard treatment (e.g. daily eye drops) or recently developed long-lasting implant technologies, encouraging further studies aimed at clinical translation.[23]

## MATERIALS & METHODS

### Animals and Anesthesia

All animal experiments were completed under the approval of the Medical College of Wisconsin’s Institutional Animal Care and Use Committee and adhere to the Association for Research in Vision and Ophthalmology (ARVO) statement for the use of animals in ophthalmic and vision research. 80 3-month-old DBA/2J mice were obtained from a commercial breeder (stock 000671, Jackson Laboratories) and housed in a 12:12 hour light/dark photoperiod with standard mouse chow and water provided *ad libitum*. For all procedures animals underwent isoflurane anesthesia in an induction chamber (VetEquip, Livermore, CA) at a rate of 5% in 100% oxygen (0.2-1 L/min) and maintained at a rate of 1-2.5% isoflurane.

### Vector Production

rAAV vectors were produced at the Medical College of Wisconsin using our previously published protocol.[47] Briefly, adherent HEK293T cells (ATCC) were triple transfected with 1) a helper plasmid expressing adenoviral derived genes required for packaging; 2) a plasmid expressing AAV Rep2 and Cap5 genes; and 3) a transgene expression plasmid (smCBA-TC40-COX2-P2A-PTGFR-TC45; herein known as CCPP). Cells were cultured in hyperflasks (Corning NY) for a period of 72 hours in DMEM + Glutamax with 2% FBS and 1% antibiotic.antimycotic (Gibco, Grand Island, NY). Following 72 hours, cells were lysed and vector purified using iodixanol gradient certification and buffer exchange, as described previously.^27^ The resulting rAAV.CCPP vector was re-suspended in HBSS+0.014% Tween20, stored at -80°C until injection, and the titer assessed using picogreen assay.^28^

### Intracameral Injection

Anesthetized animals were placed in a lateral recumbent position under an ophthalmic operating microscope (Leica Proveo 8) and the eyes dilated with 2.5% phenylephrine HCl and 1% tropicamide topical mydriatic eye drops (Akorn, Lake Forest, IL). Hydration was maintained by application of 2.5% Hypromellose eye drops (Gonak, Akorn, Lake Forest, IL, USA) to the corneal surface followed by placement of a 6mm round coverslip (Fisher Scientific, Pittsburg, PA, USA) over the eye to applanate the illumination beam and aid visualization of the anterior chamber. Rotation of the globe was stabilized using a 0.12 mm notched straight forceps (#0109025, Hagg-Streit John Weiss, UK) to secure the medial rectus muscle before a 12mm, 33-guage, sharp beveled needle (point style 2) attached to a modified 10 µL Hamilton injector syringe (Borghuis Instruments) was advanced at an oblique angle through the cornea immediately anterior to the limbus with care taken not to injure the iris, lens or posterior corneal surface. 1.5 µL rAAV vector and 0.5 µL air was injected into the aqueous humor and the pressure allowed to normalize (∼2 mins) before the needle was withdrawn, trapping an air bubble in the injection tract in order to prevent reflux.

### Slit Lamp Biomicroscopy

Mice received transillumination slit lamp exams at 7- and 10-months of age to assess for disease progression. (Topcon SL-D8Z, Topcon Medical Systems, Oakland, CA). A 2mm circular beam was aimed through the pupil and images captured using an ISO of 1200 with a Nikon camera (D810, Nikon, Japan).

### Pattern electroretinography

Mice underwent pattern electroretinography (pERG) assessments at 10-months of age (6-months post-injection) to assess global retinal ganglion cell (RGC) function. pERGs were performed in light-adapted DBA/2J mice using a Celeris pERG system (Diagnosys LLC, Cambridge, UK) with the monocular stimulator placed over the subject eye (NH, sham injected or rAAV injected) and a reference electrode placed over the contralateral eye. A bright stimulus of constant luminance (50 cd/m^2^) with high contrast (100%) square-wave profile of abruptly reversing (0.5 Hz) vertical stimuli was presented to each eye, with 800 sweeps averaged per recording in order to identify key waveform components, including N1, P1 and N2 amplitudes.

### Whole globe histology

Eyes were enucleated and placed into 4% paraformaldehyde overnight. The following day, eyes were dehydrated through a graded ethanol series, xylene cleared, and infiltrated with paraffin (Sakura Tissue Tek-VIP5; automated tissue processor). Eyes were then embedded onto tissue blocks and 4 µm sections were taken (Microm HM355s) and placed onto poly-L-lysine coated slides. Following sectioning, paraffin was removed with xylene, and sections were rehydrated and stained with hematoxylin and eosin using a Sakura Prisma automatic staining platform. Periodic acid Schiff and Masson’s trichrome staining were completed manually with standard protocols developed by Medical College of Wisconsin’s Children’s Research Institute Histology Core. Sections were imaged with an Invitrogen EVOS M5000 microscope.

### Optic nerve histology

Optic nerves were removed from enucleated globes, and the distal end was forked as to maintain orientation. Nerves were then fixed in 2% paraformaldehyde, 2% glutaraldehyde in 0.1M sodium cacodylate buffer at 4°C. Optic nerves were then post-fixed in 1% osmium tetroxide on ice and washed three times with 0.1M sodium cacodylate. Nerves were placed in a methanol series from 50-100% methanol to dehydrate. Following dehydration, optic nerves were infused with acetonitrile, followed by acetonitrile + Embed 812 (# 14120, Electron Microscopy Sciences). Nerves were placed into 100% Embed 812 overnight at room temperature, with fresh Embed 812 replaced in the morning for 6 hours. Tissue was then embedded into molds and baked overnight at 60°C. Embedded nerves were made into semi-thin sections and stained with toluidine blue. Nerves were imaged at 40x with oil immersion on a Zeiss Axio Imager.Z2 microscope and automatically tiled with Zen 3.5 blue edition software. Montaged images spanning the entire optic nerve were converted to 8-bit monochrome and processed in a semi-automated process using ImageJ (National Institutes of Health, Bethesda, MD) in order to quantify intact axons, with thresholding and quantification based on size (10-200 pixel area) and sphericity (0.25 – 1.00) to exclude areas of gliosis or swelling. In order to assess levels of gliosis and axonal swelling optic nerve images were randomly ordered into a slide deck and graded by 5 masked graders for severity based on a standard grading system. Three nerves from each treated group, and two 4-month-old control nerve images were duplicated in the analysis to allow for reproducibility to be assessed.

### Statistics

Measurement variables for IOP (pressure in mm Hg), pERG (amplitude in μV) and axon counts (absolute number/density) are continuous and sampled from normally distributed datasets. IOP was evaluated using a two-way ANOVA analysis with time and treatment as nominal variables, whilst pERG and axon counts were evaluated using one-way ANOVAs with treatment as the nominal variable; in all instances Tukey’s posttests were applied to allow for groupwise comparisons. Assessment of optic nerve gliosis and swelling yielded discontinuous nominal variables (e.g. grades) and so were compared using a Chi-squared test to evaluate whether proportions of grade severity differed across treatment groups. All comparisons were performed using Graphpad Prism 10. Consistency of optic nerve grading was visualized using Sankey plots generated using SankeyMATIC and calculated manually using weighted Cohen’s Kappa.

## DATA AVAILABILITY

All data sets are available upon reasonable request to the corresponding author

## ACKNOWLEDGEMENTS

We would like to thank Clive Wells of the MCW Electron Microscopy Core for sectioning all optic nerves and Christine Duris of the Children’s Hospital Research Institute Histology Core for their work paraffin sectioning and staining the whole globes. This work was supported through a National Eye Institute grant (R01EY032478) awarded to Dr. Lipinski. Dr. Chern was supported through a National Eye Institute Training Grant (T32EY014537).

## AUTHOR CONTRIBUTIONS

K.J.C. and D.M.L. conceptualized and designed all experiments. K.J.C., R.L.F, G.J.M. and D.M.L. performed all experiments. K.J.C. and D.M.L. drafted the manuscript. R.L.F and G.J.M. reviewed and approved the manuscript. D.M.L. supervised the project, secured all necessary regulatory approvals and provided funding.

## FIGURE LEGENDS

**Supplemental figure 1.**
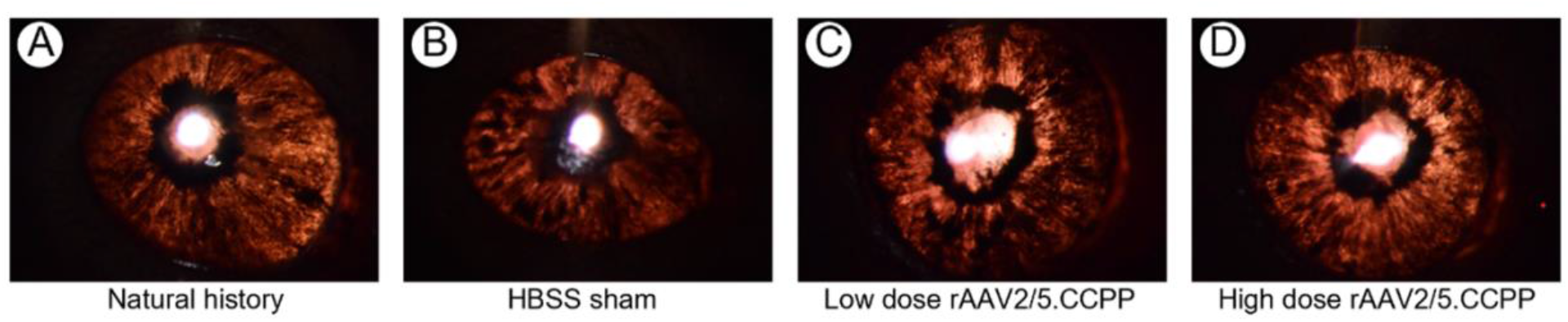
Representative transillumination images for each dosage group. Transillumination imaging acquired by slit lamp biomicroscopy and completed at the 10-month timepoint indicates normal disease progression with pigment dispersion and stromal atrophy evident in all animals at each dosage group (**A-D**).

**Supplemental figure 2.**
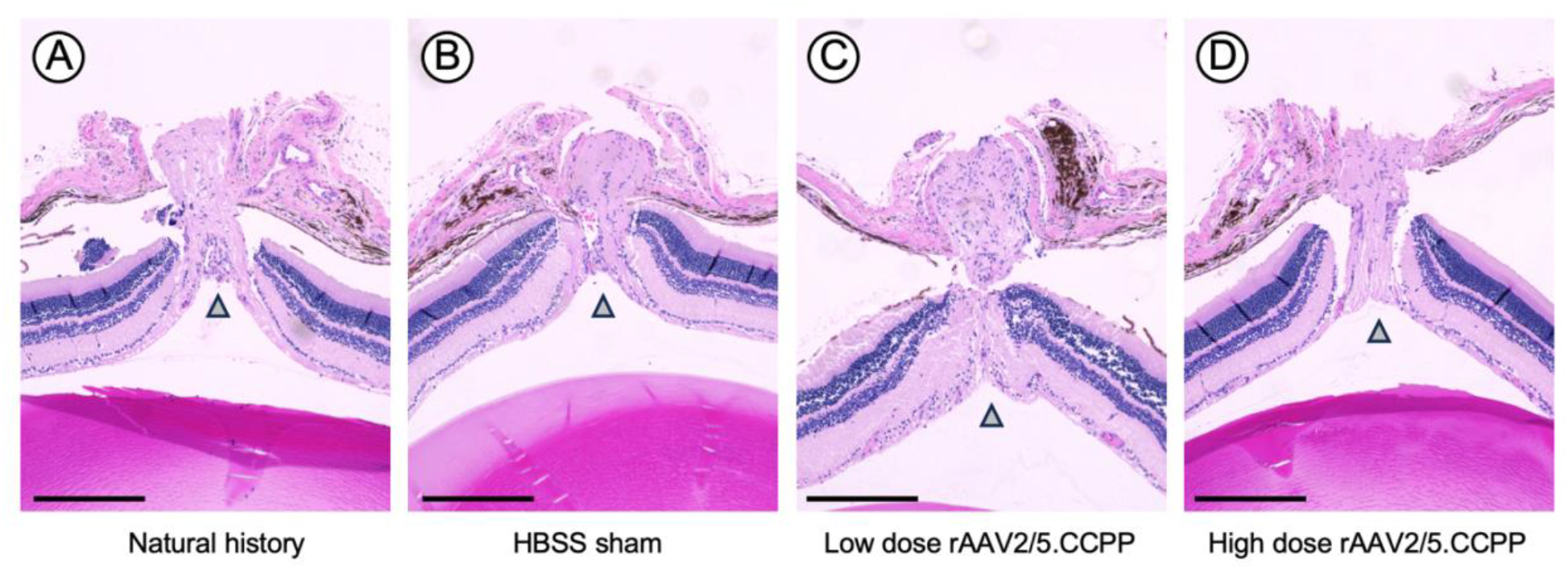
Representative coronal sections through the optic nerve head of DBA/2J mice at 10-months. Staining with hematoxylin and eosin (H&E) demonstrates more pronounced cupping of the optic nerve head in natural history (**A**) and HBSS sham treated (**B**) control eyes than in low (**C**) or high (**D**) dose rAAV2/5.CCPP injected eyes.

**Supplemental figure 3.**
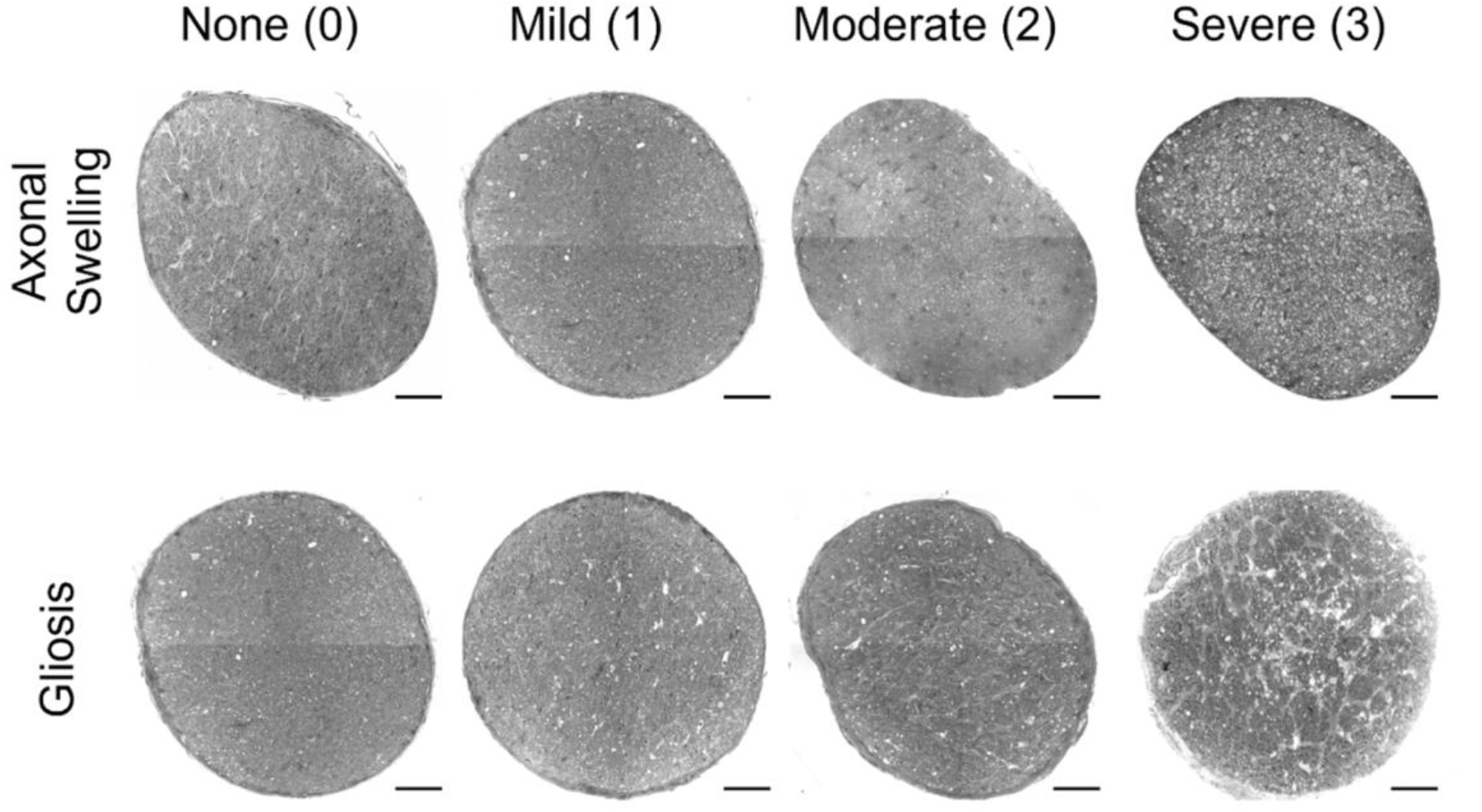
Representative images showing grade severity grading for swelling and gliosis. Representative light micrographs of toluidine blue stained coronal optic nerve sections showing examples of axonal swelling (top) and gliosis (bottom) across a range of grade severity, ranging from no damage (None or 0) through to extensive (Severe or 3). Magnification= 40x, Scale bar = 50 μm

**Supplemental figure 4.**
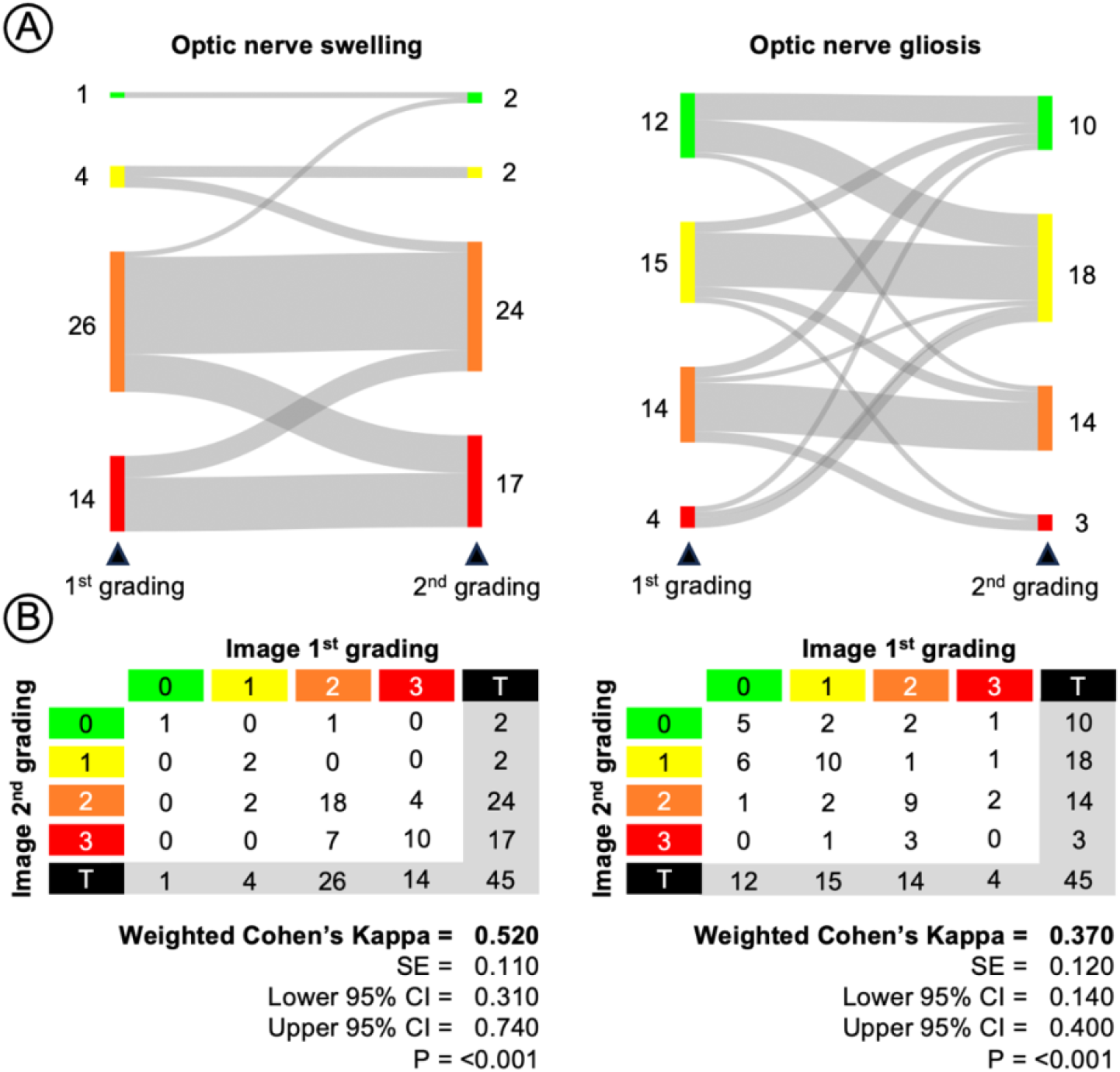
Reproducibility metrics for optic nerve grading. Sankey diagrams demonstrating consistency of scoring severity across all observers (N=5) when presented with repeated, transformed images (N=45 images total) (**A**) Weighted Cohen’s Kappa calculations for each lesion type – swelling or gliosis – based on the grading of repeated images showing a high degree of confidence (P=0.001 or <0.001) that there is only moderate (*K* range = 0.410 – 0.600) agreement between observations when grading swelling (*K* = 0.520) lesion severity, and fair (*K* range = 0.210 – 0.400) agreement between observations when grading gliosis (*K* = 0.370).

**Supplemental table 1.**
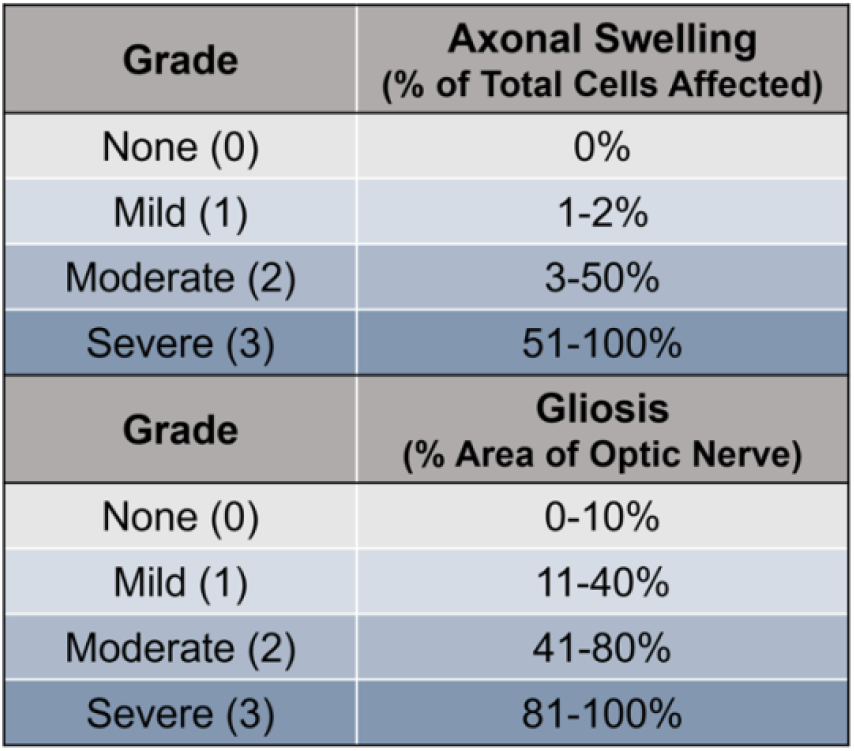
Grading scale rubric for optic nerve analysis with criteria for optic nerve analysis for axonal swelling and gliosis.

